# Chromosomal-level genome assembly of the scimitar-horned oryx: insights into diversity and demography of a species extinct in the wild

**DOI:** 10.1101/867341

**Authors:** Emily Humble, Pavel Dobrynin, Helen Senn, Justin Chuven, Alan F. Scott, David W. Mohr, Olga Dudchenko, Arina D. Omer, Zane Colaric, Erez Lieberman Aiden, David Wildt, Shireen Oliaji, Gaik Tamazian, Budhan Pukazhenthi, Rob Ogden, Klaus-Peter Koepfli

**Author notes:** Recognised as joint senior authors (Budhan Pukazhenthi, Rob Ogden, Klaus-Peter Koepfli). **Corresponding Author**: Emily Humble, Royal (Dick) School of Veterinary Studies and the Roslin Institute, University of Edinburgh, EH25 9RG, UK.

## Abstract

Captive populations provide a valuable insurance against extinctions in the wild. However, they are also vulnerable to the negative impacts of inbreeding, selection and drift. Genetic information is therefore considered a critical aspect of conservation management planning. Recent developments in sequencing technologies have the potential to improve the outcomes of management programmes however, the transfer of these approaches to applied conservation has been slow. The scimitar-horned oryx (*Oryx dammah)* is a North African antelope that has been extinct in the wild since the early 1980s and is the focus of a long-term reintroduction project. To enable the selection of suitable founder individuals, facilitate post-release monitoring and improve captive breeding management, comprehensive genomic resources are required. Here, we used 10X Chromium sequencing together with Hi-C contact mapping to develop a chromosomal-level genome assembly for the species. The resulting assembly contained 29 chromosomes with a scaffold N50 of 100.4 Mb, and displayed strong chromosomal synteny with the cattle genome. Using resequencing data from six additional individuals, we demonstrated relatively high genetic diversity in the scimitar-horned oryx compared to other mammals, despite it having experienced a strong founding event in captivity. Additionally, the level of diversity across populations varied according to management strategy. Finally, we uncovered a dynamic demographic history that coincided with periods of climate variation during the Pleistocene. Overall, our study provides a clear example of how genomic data can uncover valuable insights into captive populations and contributes important resources to guide future management decisions of an endangered species.

## Introduction

As human activities and habitat loss accelerate global species declines (Ceballos, Ehrlich, & Dirzo, 2017; Haipeng Li et al., 2016), captive and semi-captive populations are becoming increasingly important as potential sources for reintroductions (Fritz, Kramer, Hoffmann, Trobe, & Unsöld, 2017; Russell, Thorne, Oakleaf, & Ballou, 1994; Spalton, 1993). A central goal of *ex-situ* breeding programmes is therefore to achieve population viability through maintaining genetic diversity and minimising inbreeding (Frankham, Ballou, & Briscoe, 2002). Consequently, the value of genetic analysis in conservation management has long been recognised (Lacy, 1987). However, a lack of appropriate resources and baseline data has meant that in practice, genetic information is not always used. This has arguably contributed towards the failure of numerous reintroduction attempts (Robert, 2009; Tallmon, Luikart, & Waples, 2004; Weeks et al., 2011). Continued advances in sequencing technology have now made it possible to generate high resolution genomic data for practically any species, and the wider uptake of these approaches by the conservation community would undoubtedly increase the chance of successful management outcomes (Allendorf, Hohenlohe, & Luikart, 2010; Shafer et al., 2015; Supple & Shapiro, 2018; Wildt et al., 2019).

The advent of next-generation sequencing over the past decade has meant that reference genomes are now available for hundreds of species (Koepfli, Paten, Genome 10K Community of Scientists, & O’Brien, 2015). However, most genomes have been assembled using short-read sequencing technologies and as a result are highly fragmented into hundreds or thousands of scaffolds, often without any chromosomal assignment (Bradnam et al., 2013; Salzberg & Yorke, 2005). Consequently, there has been growing interest in sequencing technologies that incorporate long-range, chromosomal information to improve contiguity, reduce error rates and make downstream annotation more reliable (van Dijk, Jaszczyszyn, Naquin, & Thermes, 2018). For example, 10X Chromium sequencing uses Linked-Reads to provide long-range information, whilst Hi-C contact mapping uses structural information to build chromosome-length scaffolds (Dudchenko et al., 2017). These approaches show great promise for studies of threatened species where well characterised genomes are rarely available. Reference assemblies can aid in the development of SNP arrays, which provide a powerful approach for genotyping low quality samples (Carroll et al., 2018), whilst structural and annotation information provide the opportunity to elucidate the genetic basis of inbreeding depression, hybrid sterility and adaptation to captivity (Allendorf et al., 2010; M Kardos, Taylor, Ellegren, Luikart, & Allendorf, 2016; Knief et al., 2016).

Alongside these developments in genome assembly, whole genome resequencing is increasingly being employed to generate high resolution datasets of mapped genomic markers (Dobrynin et al., 2015; Ekblom et al., 2018; Marty Kardos, Qvarnström, & Ellegren, 2017; Robinson et al., 2016; Westbury, Petersen, Garde, Heide-Jørgensen, & Lorenzen, 2019). This has opened up the opportunity for precisely measuring genetic diversity, a critical aspect of conservation management, particularly when selecting founders for reintroduction (IUCN/SSC, 2013). However, only a handful of studies have employed genomic approaches for measuring diversity in captive species (Çilingir et al., 2019; Robinson et al., 2019; Willoughby, Ivy, Lacy, Doyle, & DeWoody, 2017) and therefore most estimates are based on traditional markers such as microsatellites. These can be associated with high sampling variance and ascertainment bias (Väli, Einarsson, Waits, & Ellegren, 2008), making comparisons across species and populations problematic. As the conservation community continues to integrate the management of captive breeding programmes and natural populations (Redford, Jensen, & Breheny, 2012), there is a growing need to reliably characterise the distribution of diversity across meta-populations.

As well as facilitating the assessment of genetic diversity, sequence data from a diploid genome assembly can be used for reconstructing demographic history. For example, studies are increasingly employing methods such as PSMC (Heng Li & Durbin, 2011)(Heng Li & Durbin, 2011) to infer past periods of population instability in wild species 08/12/2019 16:05:00 and whilst some have documented dynamic patterns that coincide with past ecological variation (Beichman et al., 2019; Mays et al., 2018), others have uncovered signals of persistent population decline (Dobrynin et al., 2015; Westbury et al., 2019). As contemporary levels of genetic diversity are largely the result of mutations and genetic drift that occurred in the past (Ellegren & Galtier, 2016), an understanding of past population dynamics can place current estimates of diversity into a historical context (Stoffel et al., 2018).

The scimitar-horned oryx (SHO), *Oryx dammah*, is a large iconic antelope and one of two mammalian species classified as extinct in the wild by the International Union for Conservation of Nature (IUCN SSC Antelope Specialist Group, 2016). The species was once widespread across North Africa, however a combination of hunting and land-use competition resulted in rapid population decline until the last remaining individuals disappeared in the 1980s (Woodfine & Gilbert, 2016). Before they were declared extinct, captive populations were established from what is thought to be around 50 individuals, mostly originating from Chad (Woodfine & Gilbert, 2016). In the decades that followed, captive SHO numbers increased to reach approximately 15,000 individuals (Gilbert, 2019). These are primarily held within unmanaged private collections such as those in the United Arab Emirates (Environment Agency of Abu Dhabi, EAD) and southern USA (Wildt et al., 2019), but also within studbook managed breeding programmes including those in Europe (European Endangered Species Program, EEP) and the USA (Species Survival Plan Program, SSP). Rapid reductions in population size, such as those associated with the founding of captive populations, are generally expected to lead to a substantial loss of genetic diversity (Frankham et al., 2002). However, an early study using mitochondrial DNA reported considerably high levels of variation in captive SHO populations (Iyengar et al., 2007). Furthermore, a recent analysis using both microsatellites and a small panel of SNPs found support for higher levels of genetic diversity in studbook managed populations, implying that diversity is not spread evenly across the globe (Ogden et al., 2020).

A programme of SHO reintroductions occurred in Tunisia between 1985–2007 (Woodfine & Gilbert, 2016) and since 2010, a large-scale effort to release the species back into its native range has been led by the Environment Agency of Abu Dhabi. To date, approximately 150 individuals have been released into Chad, and a further 350 animals are due to be reintroduced in the coming years. To enable both the selection of suitable founder individuals and effective post-release monitoring, SNP genotyping using reduced representation sequencing has been carried out across multiple populations (Ogden et al., 2020). However, to place these markers into a genomic context and improve overall resolution, more comprehensive resources are required. In this study, we used a combination of 10X Chromium sequencing and Hi-C based chromatin contact maps to generate a chromosomal-level genome assembly for the species. We additionally resequenced six individuals from across three captive populations to generate a panel of genome-wide SNPs. The resulting data were used to investigate the strength of chromosomal synteny between oryx and cattle (*Bos taurus*), elucidate patterns of diversity between mammalian species and across captive SHO populations, and reconstruct historical demography of the oryx. We hypothesised that: i) SHO and cattle would display strong chromosomal synteny given relatively recent divergence times; ii) levels of diversity in the SHO would be low compared to other mammals, considering the species is extinct in the wild; iii) intensively managed zoo populations would display higher levels of genetic diversity than largely unmanaged collections despite having smaller population sizes; and iv) patterns of past population disturbance would coincide with known periods of climatic change in North Africa.

## Materials and Methods

### Sampling and DNA extraction

Liver tissue and peripheral whole blood were collected from a male scimitar-horned oryx (international studbook #20612) from the captive herd at the National Zoological Park – Conservation Biology Institute in Front Royal, Virginia, USA. This individual represents approximately 15% of founders to the global population documented in the international studbook. Whole blood was collected into EDTA blood tubes (BD Vacutainer Blood Tube, Becton, Dickinson and Company, Franklin Lakes, NJ, USA) and stored frozen until analysis. Total genomic DNA was isolated and used to generate the *de novo* reference genome assembly (see below for details). Additional blood samples were obtained for whole genome resequencing from six individuals representing three of the main captive populations: the EEP (*n* = 2, international studbook numbers #35552 and #34412), the SSP (*n* = 2, international studbook numbers #33556 and #111029) and the EAD (*n* = 2, for further details, see Table S1). EEP blood samples were collected by qualified veterinarians during routine health procedures and protocols were approved by Marwell Wildlife Ethics Committee. Total genomic DNA was extracted between one and five times using either the Qiagen DNeasy Blood and Tissue Kit (Qiagen, Cat. No. 69504) or the QuickGene DNA Whole Blood or Tissue Kit (Kurabo Industries). Elutions were pooled and concentrated in an Eppendorf Concentrator Plus at 45°C and 1400 rpm until roughly 50 μl remained.

### 10X Genomics sequencing and assembly

Two technologies were employed to sequence and assemble the scimitar-horned oryx reference genome: 10X Genomics linked-read sequencing and chromosome conformation capture (Hi-C). For the 10X assembly, high molecular weight genomic DNA was isolated from ~2 ml of whole blood from individual #20612 using Nanobind magnetic discs (Circulomics, Inc., MD, USA). Genomic DNA concentration and purity were assessed with a Qubit 2.0 Fluorometer (ThermoFisher Scientific, MA, USA) and NanoDrop 2000 spectrophotometer (ThermoFisher Scientific, MA, USA). Capillary electrophoresis was carried out using a Fragment Analyzer (Agilent Technologies, CA, USA) to ensure that the isolated DNA had a minimum molecule length of 40 kb. Genomic DNA was diluted to ~1.2 ng/μl and libraries were prepared using Chromium Genome Reagents Kits Version 2 and the 10X Genomics Chromium Controller instrument fitted with a micro-fluidic Genome Chip (10X Genomics, CA, USA). DNA molecules were captured in Gel Bead-In-Emulsions (GEMs) and nick-translated using bead-specific unique molecular identifiers (UMIs; Chromium Genome Reagents Kit Version 2 User Guide). Size and concentration were determined using an Agilent 2100 Bioanalyzer DNA 1000 chip (Agilent Technologies, CA, USA). Libraries were then sequenced on an Illumina NovaSeq 6000 System following the manufacturer’s protocols (Illumina, CA, USA) to produce >60X read depth using paired-end 150 bp reads. The reads were assembled into phased pseudohaplotypes using Supernova Version 2.0 (10X Genomics, CA, USA). This assembly will hereafter be referred to as the 10X assembly.

### Hi-C sequencing and scaffolding

Using liver tissue from individual #20612, an *in-situ* Hi-C library was prepared as previously described (Rao et al., 2014). The Hi-C library was sequenced on a HiSeq X Platform (Illumina, CA, USA) to a coverage of 60X. The Hi-C data were aligned to the 10X Genomics linked-read assembly using Juicer (Durand et al., 2016). Hi-C genome assembly was then performed using the 3D-DNA pipeline (Dudchenko et al., 2017) and the output was reviewed using Juicebox Assembly Tools (Dudchenko et al., 2018). In cases of under-collapsed heterozygosity in the 10X assembly, one variant was chosen at random and incorporated into the 29 chromosome-length scaffolds. Alternative haplotypes are reported as unanchored sequences. This assembly will hereafter be referred to as the 10X+HiC assembly.

### Genome annotation and completeness

To identify and annotate interspersed repeat regions we used RepeatMasker v4.0.7 to screen the 10X assembly against both the Dfam_consensus (release 20170127, (Wheeler et al., 2013) and RepBase Update (release 20170127, (Bao, Kojima, & Kohany, 2015) repeat databases. Sequence comparisons were performed using RMBlastn v2.6.0+ with the -species option set to mammal. We next predicted protein-coding genes with AUGUSTUS version 3.3.2 (Stanke et al., 2006) using the gene model trained in humans. Prediction of untranslated regions was disabled and RepeatMasker repeats were provided as evidence for intergenic regions or introns. Functional annotation of the predicted genes was then performed using eggNOG-mapper v1.0.3 (Huerta-Cepas et al., 2017) against the eggNOG orthology database (Huerta-Cepas et al., 2016). The alignment algorithm DIAMOND was specified as the search tool (Buchfink, Xie, & Huson, 2015). A final set of protein-coding genes was obtained by filtering the genes predicted by AUGUSTUS for those with gene names assigned by eggNOG-mapper. Genome completeness of both the 10X and 10X+Hi-C assemblies was assessed using BUSCO v2 with 4,104 genes from the Mammalia odb9 database (Simão, Waterhouse, Ioannidis, Kriventseva, & Zdobnov, 2015) and the gVolante web interface (Nishimura, Hara, & Kuraku, 2017).

### Genome synteny

We aligned the SHO chromosomes from the 10X+HiC assembly to the cattle genome (*Bos taurus* assembly version 3.1.1, GenBank accession number GCA_000003055.5, Zimin et al., 2009) using LAST v746 (Kiełbasa, Wan, Sato, Horton, & Frith, 2011). The cattle assembly was first prepared for alignment using the command lastdb. Next, lastal and last-split commands in combination with parallel-fastq were used to align the SHO chromosomes to the cattle assembly. Coordinates for alignments over 10 Kb were extracted from the resulting multiple alignment format file and visualised using the R package RCircos v1.2.0 (Zhang, Meltzer, & Davis, 2013).

### Whole-genome resequencing and alignment

Library construction was carried out for whole genome resequencing of the six focal individuals using the Illumina TruSeq Nano High Throughout library preparation kit. Paired-end sequencing was performed on an Illumina HiSeq X Ten platform at a depth of coverage of 15X. Sequencing reads were mapped to the SHO 10X+HiC chromosomes using BWA MEM v0.7.17 (Heng Li, 2013) with the default parameters. Any unmapped reads were removed from the alignment files using SAMtools v1.9 (Heng Li, 2011). We then used Picard Tools to sort each bam file, add read groups and mark and remove duplicate reads. This resulted in a set of six filtered alignments for each of the resequenced individuals.

### SNP calling and filtering

HaplotypeCaller in GATK v3.8 (Van der Auwera et al., 2013) was first used to call variants separately for each filtered bam file. GenomicVCF files for each individual were then used as input to GenotypeGVCFs for joint genotyping. The resulting SNP dataset was filtered to include only biallelic SNPs using BCFtools v1.9 (Heng Li, 2011). We then applied a set of filters to obtain a high-quality dataset of variants using VCFtools v0.1.13 (Danecek et al., 2011). First, loci with Phred-scaled quality scores of less than 50 and genotypes with a depth of coverage less than five or greater than 38 (twice the mean sequence read depth) were removed. Second, loci with any missing data were discarded. Finally, we removed loci that did not conform to Hardy-Weinberg equilibrium with a *p*-value threshold of <0.001 and with a minor allele frequency of less than 0.16 to ensure the minor allele was observed at least twice.

### Mitochondrial genome assembly

Sequencing reads for the six resequenced individuals were mapped using BWA MEM v0.7.17 to a published mitochondrial reference genome of an SHO originating from the Paris Zoological Park (NCBI accession number: JN632677, Hassanin et al., 2012). Alignment files were filtered to contain only reads that mapped with their proper pair. Variants were called using SAMtools mpileup and BCFtools call commands and filtered to include only those with Phred quality scores over 200 using VCFtools. The resulting VCF file was manually checked and sites where the called allele was supported by fewer reads than the alternative allele were corrected. Consensus sequences for each individual were extracted using the BCFtools consensus command. We next used Geneious Prime v2019.2.1 (https://www.geneious.com) to annotate the mitochondrial consensus sequences and extract the cytochrome b, 16S and control region from each individual. Sequence similarity and haplotype frequencies were calculated using the R package pegas (Paradis, 2010). To place the mitochondrial data into a broader geographic context, the six control region sequences were aligned to 43 previously described haplotypes (NCBI accession numbers DQ159406–DQ159445 and MN689133–MN689138, Iyengar et al. 2007; Ogden et al., 2020) using Geneious Prime. A median-joining haplotype network was generated using PopArt v1.7 (Leigh & Bryant, 2015).

### Genetic diversity

We assessed genetic diversity of SHO using two genome-wide measures. First, we used VCFtools to estimate nucleotide diversity (*π*) across all six resequenced individuals based on high-quality variants called by GATK. Second, we estimated individual genome-wide heterozygosity as the proportion of polymorphic sites over the total number of sites using the site-frequency spectrum of each individual sample. For this, filtered bam files were used as input to estimate the observed folded site-frequency spectrum (SFS) using the -doSaf and -realSFS functions in the program ANGSD (Korneliussen, Albrechtsen, & Nielsen, 2014). We excluded the X chromosome and skipped any bases and reads with quality scores below 20. Genome-wide heterozygosity was then calculated as the second value of the SFS (number of heterozygous genotypes) over the total number of sites, for each chromosome separately. To compare the level of diversity in SHO with other species, we visualised genome-wide heterozygosity values for other mammalian species collected from the literature (Table S2) against census population size and International Union for Conservation of Nature (IUCN) status. Finally, assuming a per site/per generation mutation rate (μ) of 1.1×10^−08^, we used our estimate of nucleotide diversity (*π*) as a proxy for *θ* to infer long-term *N*^*e*^, given that *θ* = 4*Ne*μ.

### Demographic history

To reconstruct the historical demography of the SHO, we used the Pairwise Sequential Markovian Coalescent (PSMC, Heng Li & Durbin, 2011). This method uses the presence of heterozygous sites across a diploid genome to infer the time to the most recent common ancestor between two alleles. The inverse distribution of coalescence events is referred to as the instantaneous inverse coalescence rate (IICR) and for an unstructured and panmictic population, can be interpreted as the trajectory of *N*^*e*^ over time (Chikhi et al., 2018). To estimate the PSMC trajectory, we first generated consensus sequences for all autosomes in each of the filtered bam files from the six re-sequenced individuals using SAMtools mpileup, bcftools call and vcfutils.pl vcf2fq. Sites with a root-mean-squared mapping quality less than 30, and a depth of coverage below four or above 40 were masked as missing data. PSMC inference was then carried out using the default input parameters to generate a distribution of IICR through time for each individual. To generate a measure of uncertainty around our PSMC estimates, we ran 100 bootstrap replicates per individual. For this, consensus sequences were first split into 47 non-overlapping segments using the splitfa function in PSMC. We then randomly sampled from these, 100 times with replacement, and re-ran PSMC on the bootstrapped datasets.

To determine the extent to which the PSMC trajectory could vary, we scaled the coalescence rates and time intervals to population size and years based on three categories of neutral mutation rate and generation time. Our middle scaling values corresponded to a mutation rate of 1.1 × 10^−08^ and a generation time of 6.2 years, and were considered the most reasonable estimates for the SHO. These were based on the per site/per generation mutation rate recently estimated for gemsbok (*Oryx gazella*, Chen et al., 2019) and the generation time reported in the International Studbook for the SHO (Gilbert, 2019). Low scaling values corresponded to a mutation rate of 0.8 × 10^−08^ and a generation time of three and high scaling values corresponded to a mutation rate of 1.3 × 10^−08^ and a generation time of ten. Finally, to test the reliability of our IICR trajectories, we simulated sequence data under the inferred PSMC models and compared estimates of genome-wide heterozygosity with empirical values (Beichman, Phung, & Lohmueller, 2017). To do this, we used the program MaCS (G. K. Chen, Marjoram, & Wall, 2009) to simulate 1000 × 25 Mb sequence blocks under the full demographic model of each individual, assuming a recombination rate of 1.0 × 10^−8^ base pair per generation and a mutation rate of 1.1 × 10^−08^. Simulated heterozygosity was then calculated as the number of segregating sites over the total number of sites for each 25 Mb sequence. Empirical heterozygosity was calculated for each individual as the number of variable sites over the total number of sites in 25 Mb non-overlapping sliding windows along the genome. This was carried out using the filtered SNP dataset and the R package *windowscanr*.

## Results

### Chromosomal-level genome assembly

The genome assembly of the SHO, generated using both 10X Chromium and Hi-C technologies, had a total length of 2.7 Gb (Table 1). The use of Hi-C data successfully incorporated scaffolds into 29 chromosomes and increased the scaffold N50 by almost three-fold from 35.2 Mb to 100.4 Mb, and the contig N50 by over two-fold from 378 kb to 852 kb (Table 1). Around 149 Mb of under-collapsed heterozygosity was identified and incorporated into the assembly as unanchored sequence. The estimated GC content of the 10X-Hi-C assembly was 41.8%. BUSCO analysis of gene completeness revealed that 93.3% of core genes were complete in the 10X+Hi-C assembly which represents a marginal improvement in gene completeness compared to the 10X assembly (Table 1). Repetitive sequence content based on LTR elements, SINEs, LINEs, DNA elements, small RNAs, low complexity sequences and tandem repeats corresponded to approximately 47.63% of the genome (Table S3). SINEs and LINEs were the most common repeat elements, representing around 38% of the overall repeat content. Gene prediction using AUGUSTUS identified a total of 30,228 candidate protein-coding genes, of which 14,119 were assigned common gene names using eggNOG-mapper.

**Table 1:** Genome assembly statistics for both iterations of the SHO genome assembly. Complete core genes, complete and partial core genes, missing core genes and average number of orthologs per core gene were assessed using BUSCO v2 with the Mammalia odb9 database (4,104 genes).

|  | 10X | 10X+Hi-C |
| --- | --- | --- |
| Length (bp) | 2,720,895,635 | 2,720,101,635 |
| Scaffold N50 (bp) | 35,228,849 | 100,398,400 |
| Scaffold L50 | 21 | 11 |
| Longest scaffold (bp) | 136,126,622 | 198,955,781 |
| Contig N50 (bp) | 378,550 | 852,138 |
| GC content (%) | 41.82 | 41.83 |
| Complete core genes (%) | 92.76 | 93.25 |
| Complete & partial core genes (%) | 95.98 | 96.15 |
| Missing core genes (%) | 4.02 | 3.85 |
| Average number of orthologs per core gene | 1.05 | 1.04 |

### Genome synteny

To explore genomic synteny between SHO and cattle, we aligned the 29 chromosomes from the 10X+Hi-C assembly to the cattle assembly (BosTaurus version 3.1.1). Visualisation of the full alignment identified one chromosomal fusion between cattle chromosomes C1 and C25 which was located on SHO chromosome SHO2 (Figure 1). All remaining SHO chromosomes mapped mainly or exclusively to a single cattle chromosome, reflecting strong chromosomal synteny between the two species. Specifically, for 28 SHO chromosomes, over 90% of the total alignment length was to a single cattle chromosome, with 11 of these aligning exclusively to a single cattle chromosome.

**Figure 1:**
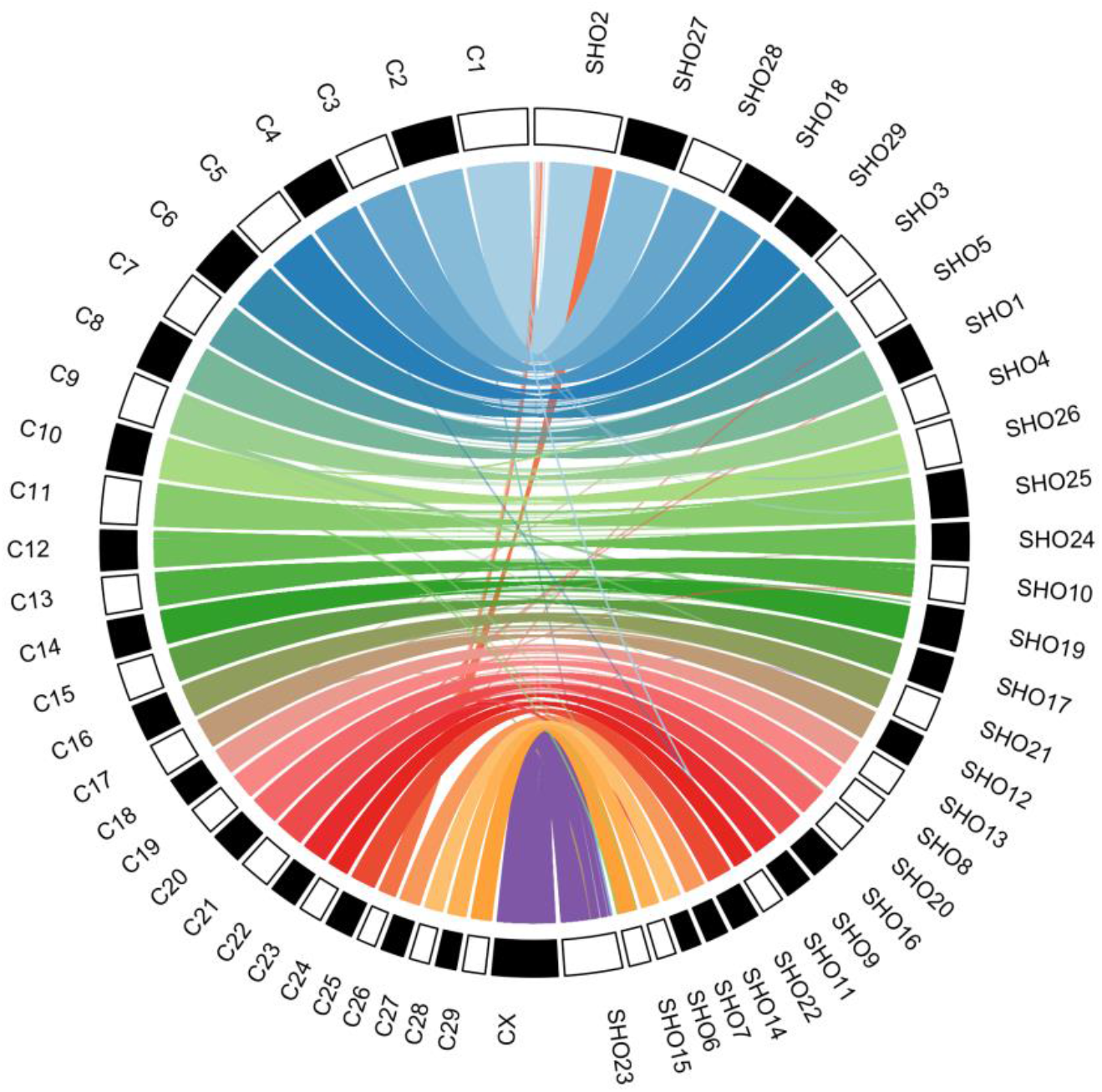
Synteny between the 29 SHO 10X+HiC chromosomes (prefixed with SHO) and the cattle chromosomes (prefixed with C). Mapping each SHO chromosome resulted in multiple alignment blocks (mean = 2.5 kb, range = 0.3 – 12.5 kb) and alignments over 10 kb are shown.

### Whole genome resequencing and SNP discovery

Whole genome resequencing of the six focal individuals resulted in an average sequencing coverage of 18.9 (min = 15.5, max = 27.2). After variant calling, a total of 12,945,559 biallelic SNPs were discovered using GATK’s best practice workflow (see Materials and Methods for details). Of these, a total of 8,063,284 polymorphic SNPs remained after quality filtering, with a mean minor allele frequency of 0.29. A full breakdown of the number of variants remaining after each filtering step is provided in Figure S1.

### Mitochondrial genome assembly

We used the whole genome resequencing data, together with a publicly available mitochondrial DNA reference sequence to assemble the mitochondrial genome for the six focal SHO individuals. An average of 1,211,796 reads per individual mapped to the reference sequence (min = 27,178, max = 5,663,594), equivalent to an average mitochondrial sequencing coverage of 3487 (min = 342, max = 7934). Across each of the six consensus sequences, a total of 125 substitutions were identified, with sequence similarity ranging between 99.5 to 100% (Table S4). Individuals from EEP and SSP breeding programmes each displayed a unique mitochondrial haplotype whilst the haplotypes of both EAD animals were identical. Furthermore, we identified a total of five control region haplotypes, five 16S haplotypes and three cytochrome b haplotypes. To place our mitochondrial data into a broader context, we compared the control region sequences for each individual with 43 previously published haplotypes. Visualization of the haplotype network revealed that all five haplotypes from this study corresponded to previously published sequences (Table S1). Haplotypes from the four EAD and SSP animals clustered together on the left-hand side of the haplotype network, whilst haplotypes from the two EEP animals clustered separately on the right-hand side of the network. This suggests that a reasonably wide proportion of the known genetic diversity for the species has been captured (Figure S2).

### Genetic diversity

Next, we investigated the level of variation in the SHO using two genome-wide measures. Our estimate for nucleotide diversity (*π*), the average number of pairwise differences between sequences, was 0.0014. Average genome-wide heterozygosity across all six individuals was in line with this, at 0.0097 (Figure 2A). Whilst this is lower than values estimated for mammals such as the brown bear and bighorn sheep, this is considerably higher than estimates for endangered species such as the baiji river dolphin and the cheetah. Furthermore, given a census population size of around 15,000 individuals, this level of diversity is in line with that of species with similar census sizes such as the orangutan and the bonobo. Among individuals, genome-wide heterozygosity ranged between 0.00076 and 0.0011, with animals from the EAD displaying the lowest levels of genome-wide heterozygosity (Figure 2B). Diversity estimates for animals from European and American captive breeding populations were similar, with American animals being slightly more diverse (Figure 2B). Genome-wide heterozygosity also varied across autosomes, with some individuals displaying larger variance in heterozygosity than others (Figure 2B). Using our estimate of genome-wide heterozygosity as a proxy for *θ*, and assuming a mutation rate of 1.1e^−8^, long-term *N*^*e*^ of the SHO was estimated to be approximately 22,237 individuals.

**Figure 2:**
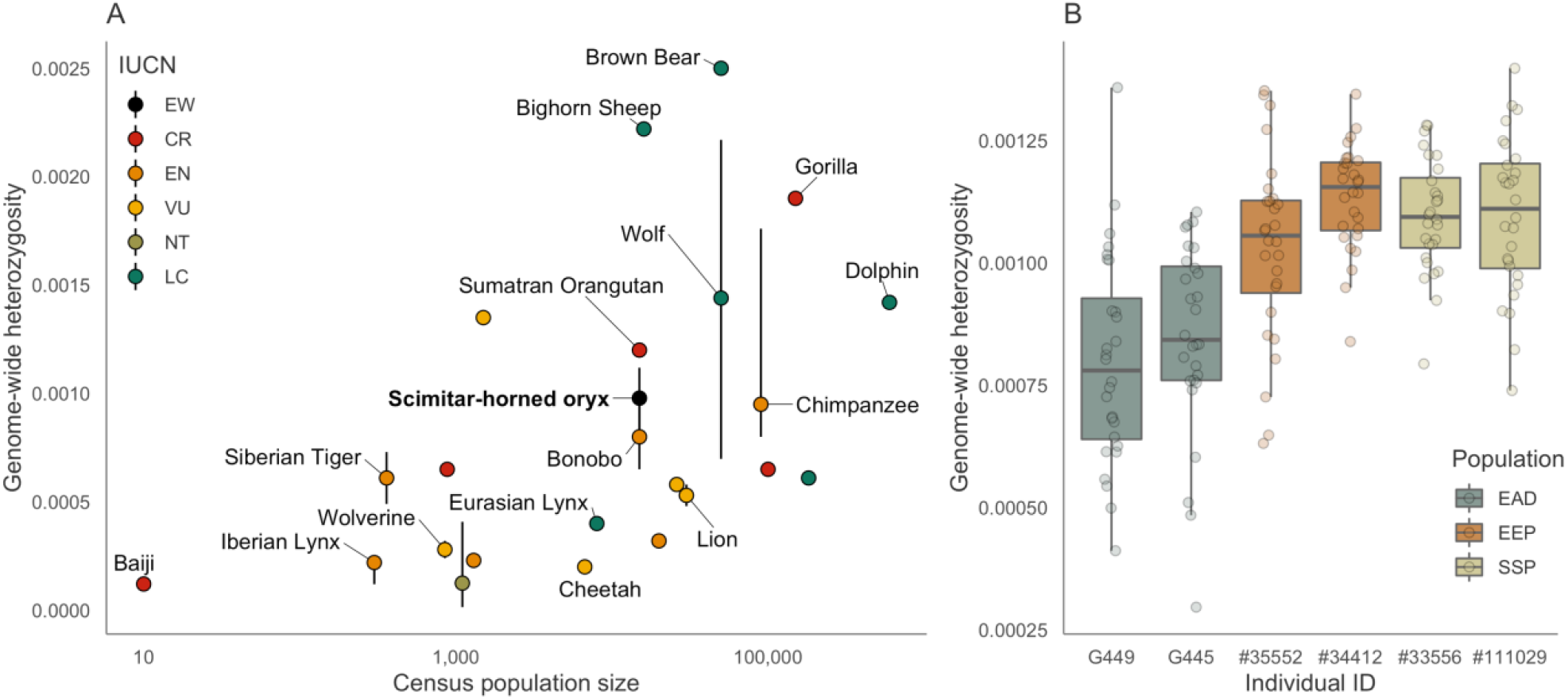
(A) Relationship between genome-wide heterozygosity and census population size for a selection of mammals, with individual points colour coded according to IUCN status. Some species names have been removed for clarity. Vertical bars correspond to the range of genome-wide heterozygosity estimates when more than one was available. For sources, see Table S2. (B) Differences in genome-wide heterozygosity across SHO individuals with colours corresponding to population. Raw data points represent the average genome-wide heterozygosity of each chromosome in each individual. Centre lines of boxplots reflect the median, bounds of the boxes reflect the 25^th^ and 75^th^ percentiles and upper and lower whiskers reflect the largest and smallest values. Further details about individual animals can be found in Table S1.

### Demographic history

To investigate historical demography of SHO, we characterised the temporal trajectory of coalescent rates using PSMC. The PSMC trajectory showed the same pattern across all six individuals and therefore the curve for only one individual (#34412 from the EEP) is presented here (Figure 3, see Figure S3 for all PSMC distributions). Assuming a generation time of 6.2 years and a mutation rate of 1.1 × 10^−8^, the trajectory could be reliably estimated from approximately 2 million years ago. It was characterised by an overall decline towards the present day, interspersed with multiple periods of elevated IICR during the Pleistocene. If IICR is assumed to be equivalent to *N*^*e*^, the period of decline during the early-mid Pleistocene reached a minimum effective population size of approximately 21,000 individuals. There was a sharp increase immediately after this, which peaked approximately 150 ka before it gradually declined again at the onset of the Last Glacial Period. After the Last Glacial Maximum 22 ka, the trajectory underwent a period of increasing IICR before estimates become unreliable. Under alternative generation and mutation rate scalings, population size and year estimates shift in either direction. For example, the peak in *N*^*e*^ around 150 ka could shift by around 15,000 individuals and by up to 70 ka. To test the reliability of our PSMC trajectories, we compared the distributions of genome-wide heterozygosity calculated from both simulated and empirical data. For all individuals, the distribution of simulated heterozygosity was highly similar to empirical values, with the average empirical heterozygosity lying within the 95% confidence intervals of the simulated distribution indicating that the PSMC models are a good fit to the data (Figure S4).

**Figure 3:**
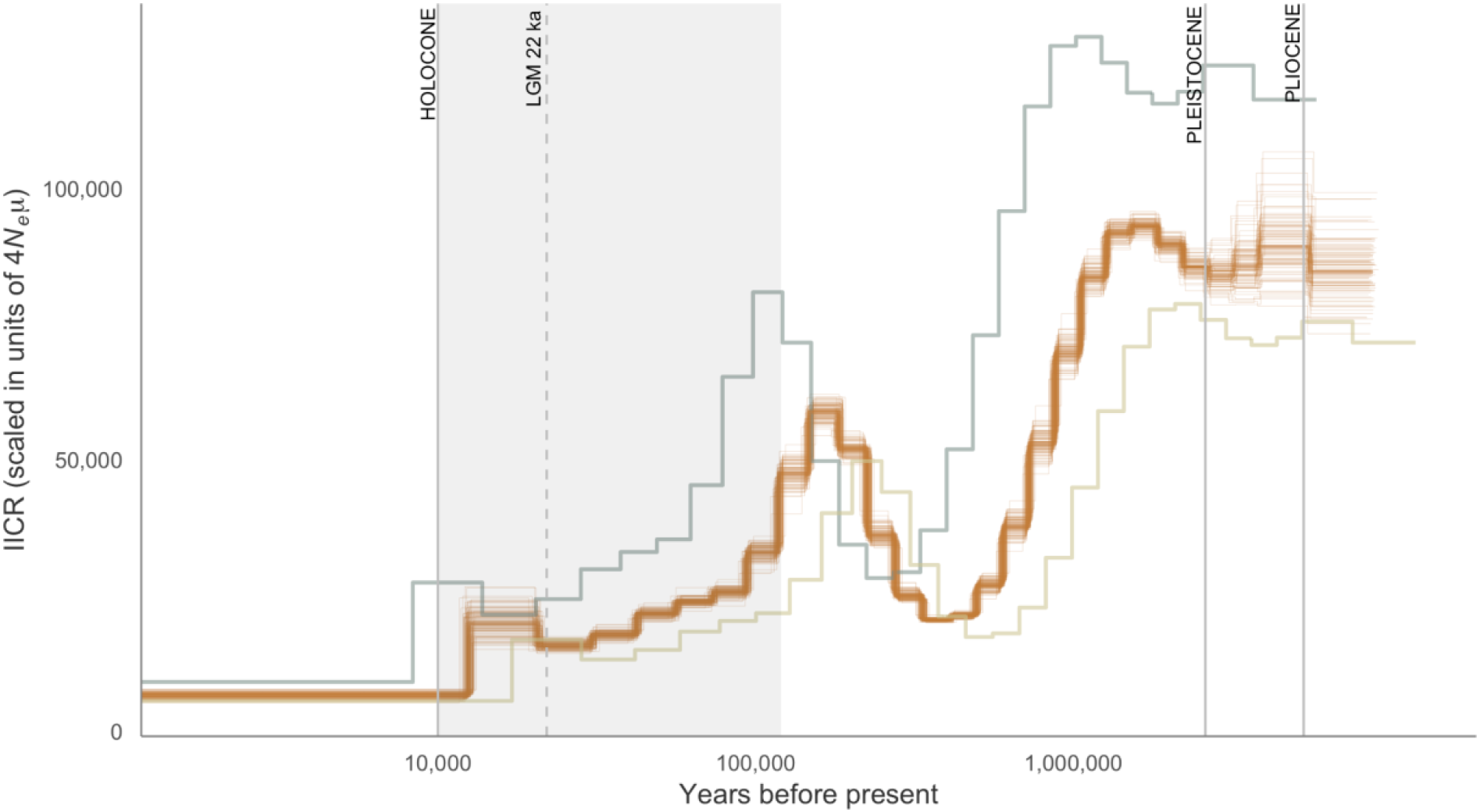
PSMC inference of the instantaneous inverse coalescent rate (IICR) through time under different scalings for SHO individual #34412 from the EEP. See Figure S3 for PSMC distributions of all individuals. The orange trajectory was scaled by a mutation rate of 1.1 × 10^−08^ and a generation time of 6.2 (medium), the grey trajectory was scaled by a mutation rate of 0.8 × 10^−08^ and a generation time of three (low) and the gold trajectory as scaled by a mutation rate of 1.3 × 10^−08^ and a generation time of 10 (high). Fine lines around the orange trajectory represent 100 bootstrap replicates. The shaded grey area corresponds to the Last Glacial Period and the Last Glacial Maximum (LGM) is indicated by the dashed line.

## Discussion

As captive populations become increasingly important for the preservation of species, it is essential that genetic resources and baseline data are available to inform population management and improve reintroduction planning. In this study, we utilised third-generation sequencing technology to generate a chromosomal-level genome assembly for the scimitar-horned oryx, a species declared extinct in the wild and the focus of a long-term reintroduction programme. We combined this with whole genome resequencing data from six individuals to characterise synteny with the cattle genome, elucidate the level and distribution of genetic diversity, and reconstruct historical demography. Our results improve our understanding of an iconic species of antelope and provide an important example of how genomic data can be used for applied conservation management.

### Genome assembly

One of the main outcomes of this study is a chromosomal-level genome assembly for the SHO, a species belonging to the subfamily Hippotraginae within the family Bovidae and superorder Cetartiodactyla. This was achieved using a combination of 10X Chromium sequencing and Hi-C contact mapping. The total assembly length was 2.7 Gb, similar to the hippotragine sable antelope (*Hippotragus niger*; Koepfli et al., 2019) and gemsbok (*Oryx gazella*; Farré et al., 2019) reference assemblies, which have total lengths of 2.9 and 3.2 Gb respectively. The use of Hi-C data successfully incorporated scaffolds into 29 chromosomes, increasing the scaffold N50 to 100.4 Mb. This is almost double that of the N50 reported for gemsbok (47 Mb, Farré et al., 2019) yet similar to that reported for the sable antelope (100.2 Mb, Koepfli et al., 2019). In contrast, the contig N50 of the 10X-Hi-C assembly was >850 kb which represents a substantial improvement over both sable antelope (45.5 kb) and gemsbok assemblies (17.2 kb). Repeat content (47.63%) was is in line with that of European bison (47.3%, Wang et al., 2017) and sable antelope assemblies (46.7%, Koepfli et al., 2019) but slightly higher than that of the Tibetan antelope (37%, Ge et al., 2013), whilst GC content was identical to that reported for the sable antelope (41.8%, Koepfli et al., 2019). Furthermore, a larger number of protein-coding genes were predicted in the SHO assembly than in studies of sable and Tibetan antelope and BUSCO analysis identified 93.3% of core genes. Our SHO assembly is therefore of very high quality and will serve as an important resource for the wider antelope and bovid research community.

### Genome synteny

To further evaluate genome completeness and to explore chromosomal synteny, we mapped the SHO chromosomes to the cattle reference genome. The resulting alignment revealed complete coverage to all chromosomes in the cattle assembly, including the X-chromosome. This is in line with the results of the BUSCO analysis and suggests that the SHO genome assembly is close to complete. Furthermore, all but one of the SHO chromosomes showed near-to, or complete chromosomal homology with cattle, indicating that the Hi-C contact mapping approach reliably anchored scaffolds into chromosomes. In general, while Bovidae genomes show a high degree of synteny, they can vary in their diploid chromosome number due to the occurrence of centric fusions (Gallagher Jr & Womack, 1992; Wurster & Benirschke, 1968). We clearly identified the fixed centric fusion between cattle chromosomes 1 and 25 that has previously been described in the oryx lineage using cytogenic approaches (Kumamoto, Charter, Kingswood, Ryder, & Gallagher, 1999). However, we found no evidence for the fusion between chromosomes 2 and 15 that has been karyotyped in some captive individuals (Kumamoto et al., 1999). Chromosomal rearrangements both within and between species have been implicated in poor reproductive performance due to the disruption of chromosomal segregation during meiosis (Hauffe & Searle, 1998; Steiner et al., 2015; Wallace, Searle, & Everett, 2002). Genotype data from additional individuals would facilitate a comprehensive assessment of structural polymorphism across captive populations of SHO using methods that utilise patterns of linkage and substructure (Knief et al., 2016).

### Genetic diversity

To assess the level of genetic diversity in the SHO we used whole genome resequencing data from six individuals originating from three captive populations. A recent meta-analysis has demonstrated that threatened species harbour reduced genetic diversity than their non-threatened counterparts due to the elevated impacts of inbreeding and genetic drift in small populations (Willoughby et al., 2015). In contrast, a handful of studies have uncovered unexpectedly high levels of diversity in species thought to have experienced strong population declines (Busch, Waser, & DeWoody, 2007; Dinerstein & McCracken, 1990; Hailer et al., 2006). While the SHO has been kept in captivity for the last 50 years, equivalent to around eight generations, it is unclear to what extent this has impacted its genetic variation. We found several lines of evidence in support for considerably high genetic diversity in the scimitar-horned oryx. First, the SHO genome assembly contained approximately 150 Mb of under-collapsed heterozygosity due to the presence of numerous alternative haplotypes. Second, we detected over 8 million high quality SNP markers, which given the small discovery pool of six individuals is relatively high for a large mammalian genome. Third, our estimates of genetic diversity were appreciably higher than in other threatened mammalian species.

These results are in some respects surprising given that the SHO underwent a period of rapid population decline in the wild, followed by a strong founding event in captivity. However, the species has bred well in captivity, reaching approximately 15,000 individuals in the space of several decades. This is likely to have reduced the strength of genetic drift, which alongside individual-based management, may have prevented the rapid loss of genetic diversity. This is in line with theoretical expectations that only very severe (i.e. a few tens of individuals) and long-lasting bottlenecks will cause a substantial reduction in genetic variation (Nei, Maruyama, & Chakraborty, 1975). With this in mind, it is also possible that the original founder population size was larger than previously thought, particularly for the EAD population, where records are generally sparse. Additionally, as contemporary levels of genetic diversity are largely determined by long-term *N*^*e*^ (Ellegren & Galtier, 2016), we cannot discount the possibility that historical patterns of abundance have contributed to the variation we see today.

Nevertheless, caution must be taken when comparing estimates of diversity across species as the total number of variable sites, and therefore genetic variation, is sensitive to SNP calling criteria (Hohenlohe et al., 2010; Shafer et al., 2017). Furthermore, there are multiple ways to measure molecular variation (Hahn, 2018). However, our results are broadly in line with similar species such as the sable antelope, where a comparable number of variants were called in a similar number of individuals (Koepfli et al., 2019). Additionally, our estimates of genome-wide heterozygosity were calculated based on genotype likelihoods and therefore should be robust to sensitivities resulting from filtering (Korneliussen et al., 2014). Finally, we took care to compare our estimates of genetic diversity with equivalent measures in the literature. Therefore, we expect our measures of genetic variation to reflect the true level of diversity in the species.

To characterize the distribution of diversity in the SHO we compared genome-wide heterozygosity among captive populations. Diversity estimates varied between groups, with animals from the EAD showing overall lower levels of diversity than those from European and American captive breeding populations. However, this comparison is based on estimates for a small number of individuals and therefore may not be a true reflection of the overall variation in genetic diversity. Nevertheless, this pattern is consistent with studies both in SHO and Arabian oryx (*Oryx leucoryx*) that found diversity to be lower in unmanaged populations than in studbook managed populations (El Alqamy, Senn, Roberts, McEwing, & Ogden, 2012) and suggests that captive breeding programmes have been successful at maintaining genetic diversity. We also observed variation in the genetic diversity of individual chromosomes, a pattern which has been demonstrated across a wide variety of taxa (Doniger et al., 2008; Nordborg et al., 2005; The International SNP Map Working Group, 2001). Chromosomal variation in heterozygosity can arise through numerous mechanisms including recombination rate variation, mutation rate variation and selection (Begun & Aquadro, 1992; Hodgkinson & Eyre-Walker, 2011; Martin et al., 2016) and further studies will be required to understand the biological significance of these patterns in more detail.

### Historical demography

To provide insights into the historical demography of the SHO, we quantified the trajectory of coalescence rates using PSMC. This method does not necessarily provide a literal representation of past population size change as it assumes a panmictic Wright-Fisher population (Mazet, Rodríguez, Grusea, Boitard, & Chikhi, 2016). Nevertheless, fluctuations in the trajectory provide insights into periods of past population instability which may be attributed to factors including population decline, population structure, gene flow and selection (Beichman et al., 2017; Chikhi et al., 2018; Mazet et al., 2016; Schrider, Shanku, & Kern, 2016). The PSMC trajectory of the SHO was characterised by an initial expansion approximately 2 million years ago which coincides with the appearance of present day bovid tribes in the fossil record (Bibi, 2013). This was followed by periods of disturbance during the mid-Pleistocene and at the onset of the Last Glacial Period, although these time points shift in either direction under alternative scalings. Similar PSMC trajectories have been observed in other African grassland species such as the gemsbok, greater kudu and impala (L. Chen et al., 2019). Climatic variability in North Africa during these time periods was associated with repeated expansion and contraction of suitable grassland habitat (Dupont, 2011), which is likely to have driven population decline or fragmentation in the SHO. This is consistent with previous findings that ecological variation associated with Pleistocene climate change has shaped the population size and distribution of ungulates in Africa (Lorenzen, Heller, & Siegismund, 2012).

Interestingly, despite the expansion of suitable SHO habitat after the Last Glacial Maxima, the PSMC trajectory does not return to historic levels. PSMC has little power to detect demographic change less than 10,000 years ago (Heng Li & Durbin, 2011), however it is possible that increased human activities during this time-period impacted population numbers. This is in line with a recent study that attributed widespread declines in ruminant populations during the late Pleistocene to increasing human effective population size (L. Chen et al., 2019). Sequencing data from additional individuals will facilitate the reliable estimation of recent population size parameters using either site-frequency based methods or approximate Bayesian computation (Excoffier, Dupanloup, Huerta-Sánchez, Sousa, & Foll, 2013; Pujolar, Dalén, Hansen, & Madsen, 2017; Stoffel et al., 2018).

### Implications for management

The outcome of this study provides important information for selecting source populations for reintroduction. In particular, our assessment of genetic diversity indicates that founders from the EAD should be supplemented with individuals from recognised captive breeding programmes. This would serve to maximise the representation of current global variation and increase the adaptive potential of release herds. Furthermore, our chromosomal genome assembly will provide a reference for generating mapped genomic markers in additional individuals and for developing complementary genetic resources such as genotyping arrays (Wildt et al., 2019). This will facilitate detailed individual-based studies into inbreeding, relatedness and admixture that will help improve breeding recommendations and hybrid assessment as well as enable post-release monitoring. Moreover, access to genome annotations will open up the opportunity for identifying loci associated with functional adaptation in both the wild and captivity. Overall, these approaches will contribute towards an integrated global management strategy for the scimitar-horned oryx and support the transfer of genomics into applied conservation.

## Conclusions

We have generated a chromosomal-level genome assembly and used whole genome resequencing to provide insights into both the contemporary and historical population of an iconic species of antelope. We uncovered relatively high levels of genetic diversity and a dynamic demographic history, punctuated by periods of large effective population size. These insights provide support for the notion that only very extreme and long-lasting bottlenecks lead to substantially reduced levels of genetic diversity. At the population level, we characterised differences in genetic variation between captive and semi-captive collections that emphasise the importance of meta-population management for maintaining genetic diversity in the remaining populations of scimitar-horned oryx.

## Data accessibility

The 10X Chromium sequencing reads are available at XXXX. The scimitar-horned oryx Hi-C assembly is available on the DNA ZOO website (www.dnazoo.org/assemblies/Oryx_dammah). Whole genome resequencing data have been deposited on the European Nucleotide Archive (accession number XXXX). Mitochondrial control region, cytochrome b and 16S mitochondrial haplotypes have been deposited on NCBI under accession number XXXX. Code for the analysis of resequencing data is available at https://github.com/elhumble/oryx_reseq.

## Author contributions

KPK, RO, HS, BP & EH conceived and designed the study. AFS and DWM carried out the 10X Chromium genome sequencing and assembly. OD, ADO, ZC and ELA carried out Hi-C genome sequencing and assembly. JC and BP contributed materials and funding. PD carried out BUSCO analysis and genome annotation with input from GT. SO contributed to mitogenome assembly and analysis. EH analysed the whole genome resequencing data and wrote the manuscript. All authors commented on and approved the final manuscript.

## Acknowledgements

We would like to thank the EAD and all EAZA and AZA SSP institutions that provided samples for this study. We would also like to acknowledge Tania Gilbert at Marwell Wildlife for advice and for access to the international studbook. ELA was supported by an NSF Physics Frontiers Center Award (PHY1427654), the Welch Foundation (Q-1866), a USDA Agriculture and Food Research Initiative Grant (2017-05741), an NIH 4D Nucleome Grant (U01HL130010), and an NIH Encyclopedia of DNA Elements Mapping Center Award (UM1HG009375). Whole-genome resequencing was carried out by Edinburgh Genomics.

